# RNA-binding protein altered expression and mislocalization in multiple sclerosis

**DOI:** 10.1101/829457

**Authors:** Katsuhisa Masaki, Yoshifumi Sonobe, Ghanashyam Ghadge, Peter Pytel, Paula Lépine, Florian Pernin, Qiao-Ling Cui, Jack P. Antel, Stephanie Zandee, Alexandre Prat, Raymond P. Roos

## Abstract

**Objective:** Nuclear depletion and mislocalization of RNA-binding proteins (RBPs) trans-activation response DNA-binding protein of 43 kDa (TDP-43) and fused in sarcoma (FUS) are thought to contribute to the pathogenesis of a number of disorders, including amyotrophic lateral sclerosis (ALS). We recently found that TDP-43 as well as polypyrimidine tract binding protein (PTB) have decreased expression and mislocalization in oligodendrocytes in demyelinated lesions in an experimental mouse model of multiple sclerosis (MS) caused by Theiler’s murine encephalomyelitis virus infection.

**Methods:** The latter finding prompted us to investigate TDP-43, FUS, and PTB in the demyelinated lesions of MS and in *in vitro cultured* human brain-derived oligodendrocytes.

**Results:** We found: i) mislocalized TDP-43 in oligodendrocytes in active lesions in some MS patients; ii) decreased PTB1 expression in oligodendrocytes in mixed active/inactive demyelinating lesions; iii) decreased nuclear expression of PTB2 in neurons in cortical demyelinating lesions; iv) nuclear depletion of TDP-43 in oligodendrocytes under metabolic stress induced by low glucose/low nutrient conditions compared to optimal culture conditions.

**Conclusion:** TDP-43 has been found to have a key role in oligodendrocyte function and viability, while PTB is important in neuronal differentiation, suggesting that altered expression and mislocalization of these RBPs in MS lesions may contribute to the pathogenesis of demyelination and neurodegeneration. Our findings also identify nucleocytoplasmic transport as a target for treatment.

## Introduction

Multiple sclerosis (MS) is a chronic inflammatory disease of the central nervous system (CNS) characterized pathologically by sharply demarcated plaques of demyelination along with axonal injury and reactive astrocytosis. Clinically MS often runs a relapsing-remitting course that is reflected in different histopathologic patterns of demyelinating plaques, which are typically described as active, mixed active/inactive or “chronic” ^1^. Progressive clinical disability is linked to demyelination in the white matter and cerebral cortex as well as cortical atrophy. The etiology and pathological mechanisms that drive white matter demyelination and neurodegeneration remain poorly understood.

Here we identify abnormalities in the expression and localization of RNA-binding proteins (RBPs) in MS lesions and in oligodendrocytes exposed *in vitro* to metabolic stress conditions. RBPs bind to double or single stranded RNA in cells and participate in the formation of ribonucleoprotein complexes (reviewed in ^2^). These proteins play a key role in RNA biology, such as splicing, polyadenylation, mRNA stabilization, mRNA localization, and translation. Most RBPs reside in the nucleus in order to carry out their function, and shuttle across the nuclear membrane in association with mRNAs. Decreased expression and mislocalization of RBPs, including trans-activation response (TAR) DNA-binding protein of 43 kDa (TDP-43) and fused in sarcoma (FUS), have been implicated in the pathogenesis of amyotrophic lateral sclerosis (ALS) and frontotemporal dementia ^2^.

We recently investigated the expression and mislocalization of RBPs in infections with Theiler’s murine encephalomyelitis virus (TMEV), a member of the *Cardiovirus* genus of *Picornaviridae* ^3^. Polypyrimidine tract binding protein (PTB) is known to have an abnormal nuclear localization in TMEV-infected cells as a result of interference of nucleocytoplasmic transport by a TMEV protein (reviewed in ^4^); the mislocalization has been hypothesized to play a role in neuronal dysfunction in TMEV-induced disease ^5^. We questioned whether other RBPs were also abnormally distributed in TMEV infections, and might contribute to disease pathogenesis. We found that TDP-43 and FUS, in addition to PTB, were mislocalized in TMEV-infected neural cells, including oligodendrocytes, in demyelinating lesions. Since TMEV-induced immune-mediated demyelination serves as an experimental model of MS, we carried out the present investigation in order to determine whether the expression pattern and localization of these RBPs were also abnormal in MS. We focused on three RBPs - TDP-43, FUS, and PTB (which included paralogs PTB1 and PTB2) - since they have been found to play an important role in TMEV infections and/or ALS.

In the present study, we found a number of examples of abnormal expression and mislocalization of RBPs in MS lesions and in *in vitro* cultured oligodendrocytes: i) mislocalization of TDP-43 in oligodendrocytes in active demyelinating lesions; ii) decreased expression of PTB1 in oligodendrocytes in mixed active/inactive demyelinating lesions; iii) decreased nuclear expression of PTB2 in neurons in cortical demyelinating lesions; iv) nuclear depletion of TDP-43 in cultured oligodendrocytes under metabolic stress conditions. These results suggest that altered expression of RBPs in demyelinating lesions of MS may disrupt key cellular processes, and thereby contribute to the pathogenesis of demyelination and neurodegeneration. A recent publication demonstrated that TDP-43 is important for oligodendrocyte survival and myelination ^6^. In addition, PTB is known to be critically important for splicing repression and neuronal differentiation ^7, 8^. The significance of these two RBPs to central nervous system function emphasizes the relevance of our results, and the potential importance of nucleocytoplasmic transport as a target for treatment.

## Methods

**Ethics statement, Human Samples, Tissue preparation and immunohistochemistry/immunofluorescence (see Supplement)**

### Staging of demyelinating lesions

We classified MS plaques into three stages ^1^ based on the density of macrophages: i) active - lesions densely and diffusely infiltrated with macrophages, ii) mixed active/inactive - lesions with macrophages restricted to the periphery, and iii) inactive – lesions with no increase in macrophage numbers within the plaque. We classified cortical plaques into three subtypes: leukocortical (involving both white matter and cortex), intracortical, and subpial (superficial cortical).

### Semi-quantitative analysis of RBPs mislocalization

Sections from blocks of cerebral cortex and white matter in all MS cases were stained with DAB or by fluorescence for RBPs that included TDP-43, PTB1, PTB2 and FUS. A semi-quantitative assessment of RBP nuclear depletion and mislocalization or decreased expression in demyelinating lesions was performed by taking digital photographs with a CMOS camera at a resolution of 1636 × 1088 pixels with a ×20 (0.75 NA) objective. At least 3 different photographs of areas of one lesion that were more than 1 mm apart from each other in x and y directions were randomly taken for every demyelinating lesion. At least 100 neuronal or glial cells per each area were identified on the basis of cytological features^11^ and scored based on the degree of mislocalization or decreased nuclear expression of RBPs compared to normal appearing white matter from the same case stained at the same time: -no or minimal; + mild (10–30 cells); ++ moderate (30–100 cells); +++ cases prominent (>100 cells (see Table e-1).

### *In vitro* studies

Oligodendrocytes were isolated and cultured from samples of 4 surgical resections (3 adults cases: 2 males, ages 57 and 38 and 1 female, age 51 and one pediatric case: male, age 7) that did not involve malignancies, as previously described ^12, 13^. The tissue was from a site distant from visible pathology. The isolation technique involved initial dissociation of tissue using trypsin digestion followed by Pespan style=“font-family:Arial”> gradient centrifugation to remove myelin. The total cell fraction was plated onto a non-coated flask that was kept overnight at 37°C to allow adhesion of the microglia fraction. Floating cells were then recovered (>90% were O4^+^) and plated into 12 well poly-L-lysine and extracellular matrix-coated chamber slides (30,000 cells per well) in defined medium (referred to as N1) consisting of Dulbecco’s modified essential medium DMEM-F12 (Sigma-Aldrich) supplemented with N1 (Sigma-Aldrich), 0.01% bovine serum albumin,1% penicillin–streptomycin, B27 (Invitrogen), platelet-dreived growth factor with two A subunits (10 ng/ml), basic fibroblast growth factor (10 ng/ml), and triiodothyronine (2 nM) (Sigma-Aldrich). After 4 days, media in individuor with DMEM with 0.25g/L glucose (referred to as low glucose (LG). After an additional 2 or 6 days of N1 or LG treatment, cells were incubated with monoclonal O4 antibody and propidium iodide (PI) for 15 min at 37°C. Cells were fixed with 4% paraformaldehyde for 10 minutes at room temperature and then stained with a secondary antibody directed against O4, goat anti-mouse IgM conjugated to AF-647 (SouthernBiotech). After permeabilization buffer with 0.1% Triton X-100, the cells were stained with polyclonal anti-TDP-43 antibody (Proteintech) for 1 hour at 37°C followed by goat anti-rabbit polyclonal antibody conjugated to AF-488 and Hoescht 33258 (Invitrogen) for 30 minutes. Cells were then examined using an epifluorescent microscope (Zeiss, Oberkochen, Germany) to determine the percent of O4 cells that were PI^+^ cells and % of cells that showed predominantly nuclear versus cytoplasmic distribution of TDP-43. Data were derived by blinded observers counting 75-100 cells per condition. Data between LG and N1 conditions were compared using a paired t-test.

### Statistical analysis

Data were analyzed using GraphPad Prism version 7.0a and are expressed as means ± standard error of the mean. Significance was assessed using Student’s t test, and P values less than 0.05 were considered statistically significant.

### Data availability statement

Any data not published within the article will be shared by request from any qualified investigator in anonymized form.

## Results

### TDP-43 in ALS

In ALS, TDP-43 is depleted from the nucleus in some motor neurons, and localized in aggregates in the cytoplasm (Fig 1A, arrows), while other neurons and glial cells have TDP-43 in its normal location in the nucleus (Fig 1A, B, arrowhead), At times phosphorylated TDP-43 (pTDP-43) is present in the cytoplasmic aggregates (Fig 1C) Two other RBPs, PTB2 and FUS, maintain their normal nuclear localization in cells in the same region that have TDP-43 mislocalization (Fig. 1D, F). PTB1 was not detected in motor neurons (Fig. 1E), since it is known to have a limited distribution in this cell type ^14^. In contrast to these finding in ALS, a predominant nuclear localization of TDP-43 is present in neurons and oligodendrocytes in human control CNS tissue (Fig. 1G, H).

**Figure 1.**
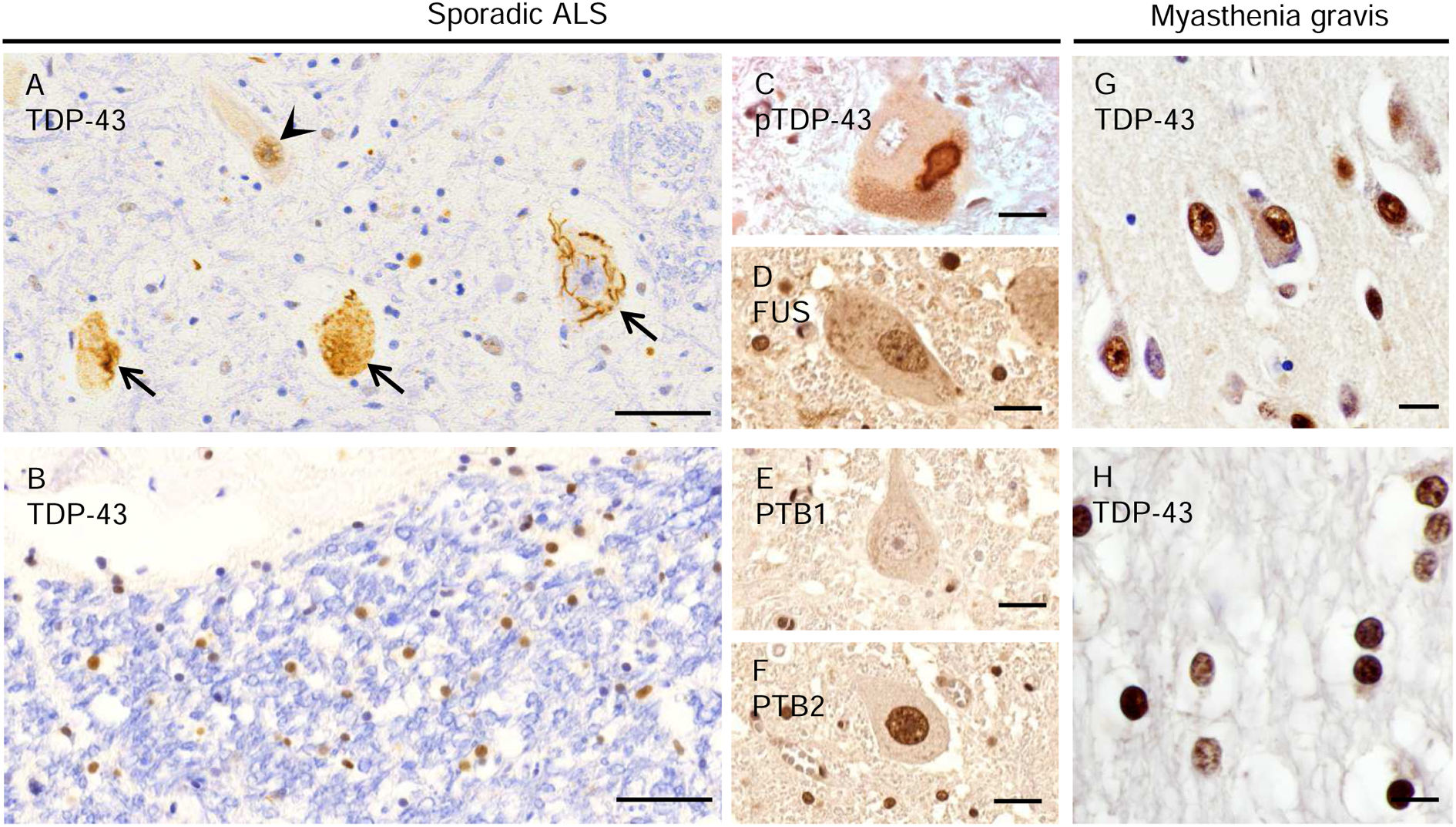
Expression pattern of RBPs in ALS and a control patient. (A-F) A case of sporadic ALS. (A) TDP-43 is expressed normally in the nucleus of some spinal cord motor neurons (*arrowhead)*, but depleted from the nucleus of affected motor neurons, forming aggregates (*large arrows*). (B) TDP-43 is localized in the nuclei of glial cells in spinal cord white matter. (C) A cytoplasmic inclusion in spinal cord motor neurons contains phosphorylated TDP-43 (pTDP-43). As expected in normal motor neurons, (D) FUS is present in the nucleus, (E) PTB1 is not present, and (F) PTB2 is present in the nucleus. (G, H) A control patient with myasthenia gravis. Expression of TDP-43 is mainly seen in the nuclei of cortical neurons (G) and white matter oligodendrocytes (H). Scale bars: 50 µm (A, B), 20 µm (C-G) and 10 µm (H).

### Altered localization and expression of RBPs in oligodendrocytes in white matter plaques

TDP-43 was mislocalized to the cytoplasm in glial cells in active demyelinating lesions from patients MS#3 and 13 to a moderate degree (Table e-1) (MS#3 - Fig. 2A-H). Double immunofluorescence demonstrated that this mislocalization was present in CNPase-positive oligodendrocytes to a significant extent (Fig. 2I, J); the nuclear depletion and cytoplasmic mislocalization was statistically significantly greater (P = 0.0003) when compared with oligodendrocytes in the periplaque white matter (PPWM) (Fig. 2I, J). Similar findings were also present in all three active demyelinating plaques in the case of MS#13 (Fig. e-1). The TDP-43 mislocalization was not a result of cellular death since the cell and its nucleus had normal morphology and there was no evidence of cleaved caspase-3 staining. Although the oligodendrocytes in active plaques in the CNS tissue from MS#3 and 13 exhibited TDP-43 mislocalization, this was not the case with the active plaques from a biopsy of a tumefactive MS lesion in MS#14 and from another MS case with three active plaques. No abnormalities were found with respect to the normal nuclear localization and expression of FUS in active plaques.

**Figure 2.**
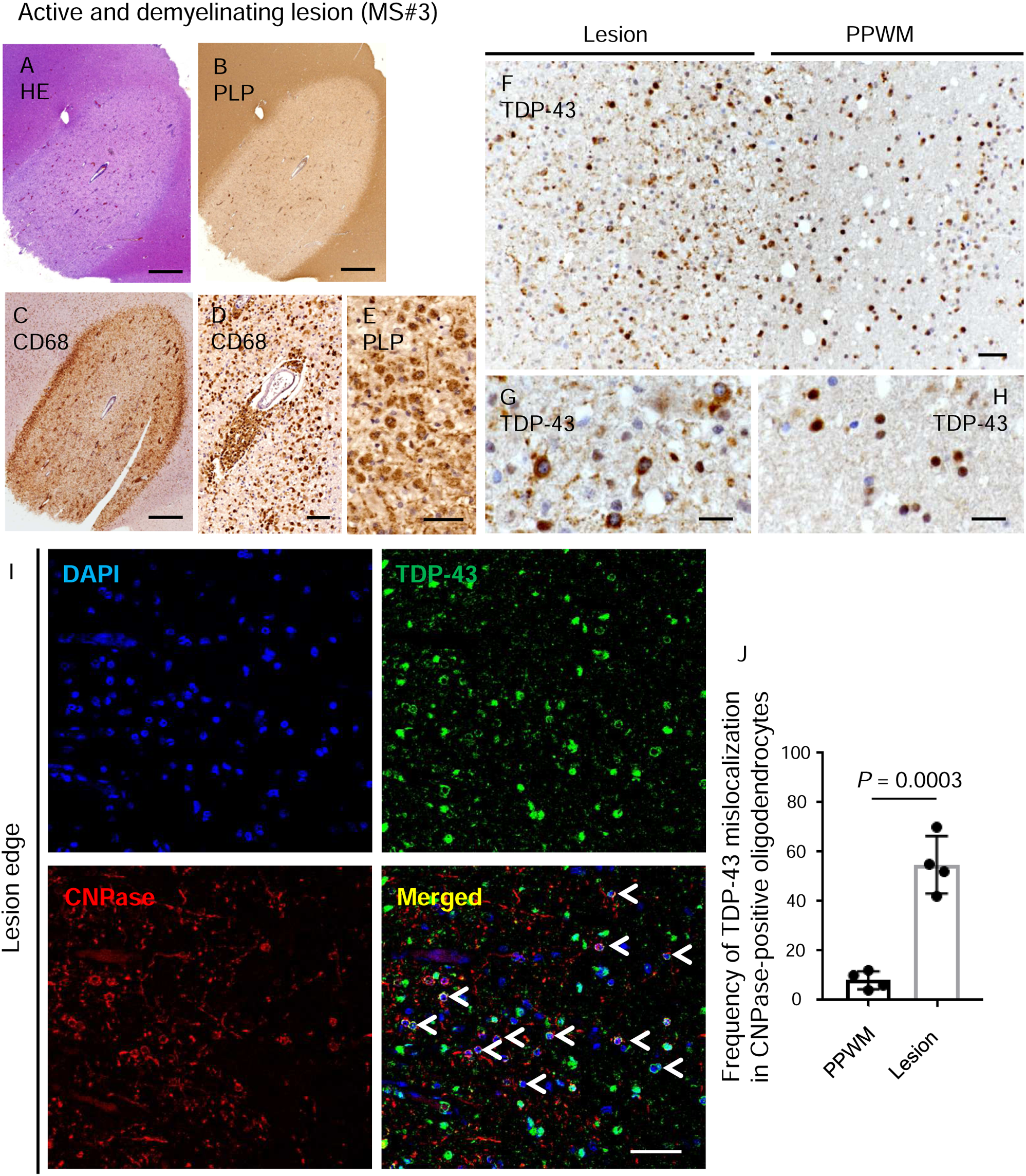
Altered expression of PTB1 and TDP-43 in oligodendrocytes of MS plaques. (A-G) An active and demyelinating lesion from MS#3. (A) HE staining shows perivascular inflammation with mononuclear cell infiltration. (B, C) Immunostaining for PLP and CD68 identifies a marked loss of myelin with dense infiltration of macrophages. (D) Numerous CD68-positive macrophages are seen in the perivascular area and parenchyma of the lesion center. (E) Foamy macrophages phagocytosing PLP-positive myelin debris. (F-H) Boundary of active demyelinating lesion of MS#3. (F, H) TDP-43 is localized in the nuclei of glial cells in peri-plaque white matter (PPWM). (F, G) TDP-43 is mislocalized to the cytoplasm of glial cells in a demyelinating lesion. (I) Double immunofluorescence shows mislocalization of TDP-43 in CNPase-positive oligodendrocytes at the edge of a demyelinating plaque (H, *arrowheads*). (J) TDP-43 mislocalization in CNPase-positive oligodendrocytes. TDP-43 mislocalization in oligodendrocytes is significantly greater in demyelinating lesions than PPWM. Each dot for this quantitation and subsequent ones represents a photograph from a separate region of the plaque. Scale bars: 1 mm (A-C), 50 µm (D-F, I) and 20 µm (G, H).

In addition to our finding of mislocalization of TDP-43 in some active plaques, there was decreased expression of PTB1 in oligodendrocytes in mixed active/inactive demyelinating lesions (MS# 2, 6, 10-13, Table e-1); however, cytoplasmic mislocalization of PTB1 was not seen in these lesions. In addition, TDP-43 and FUS were present in the nucleus in mixed active/inactive demyelinating lesions. The decreased expression of PTB1 ranged from mild to prominent (Table e-1). Although PTB1 had its expected nuclear staining in normal appearing white matter (NAWM) in the case of MS#10 (Fig. 3B, E), there was markedly decreased expression in the nuclei and cytoplasm of CNPase-positive oligodendroglia in demyelinated and partly remyelinated lesions (Fig. 3C, D, F, G); this lesion had macrophages present in the periphery as typical of active-inactive plaques. The decrease in expression was statistically significant compared to oligodendrocytes in the PPWM (P = 0.0007) (Fig. 3H). Cells with decreased PTB1 expression had normal morphology and there was no evidence of caspase-3 staining. In the case of MS#6, PTB1 expression was also diminished in mixed active/inactive demyelinating lesion (Fig. 3I-K); again, macrophages were present in the periphery of this plaque. The decrease in expression was statistically significant compared to oligodendrocytes in the PPWM (P = 0.0003) (Fig. 3K).

**Figure 3.**
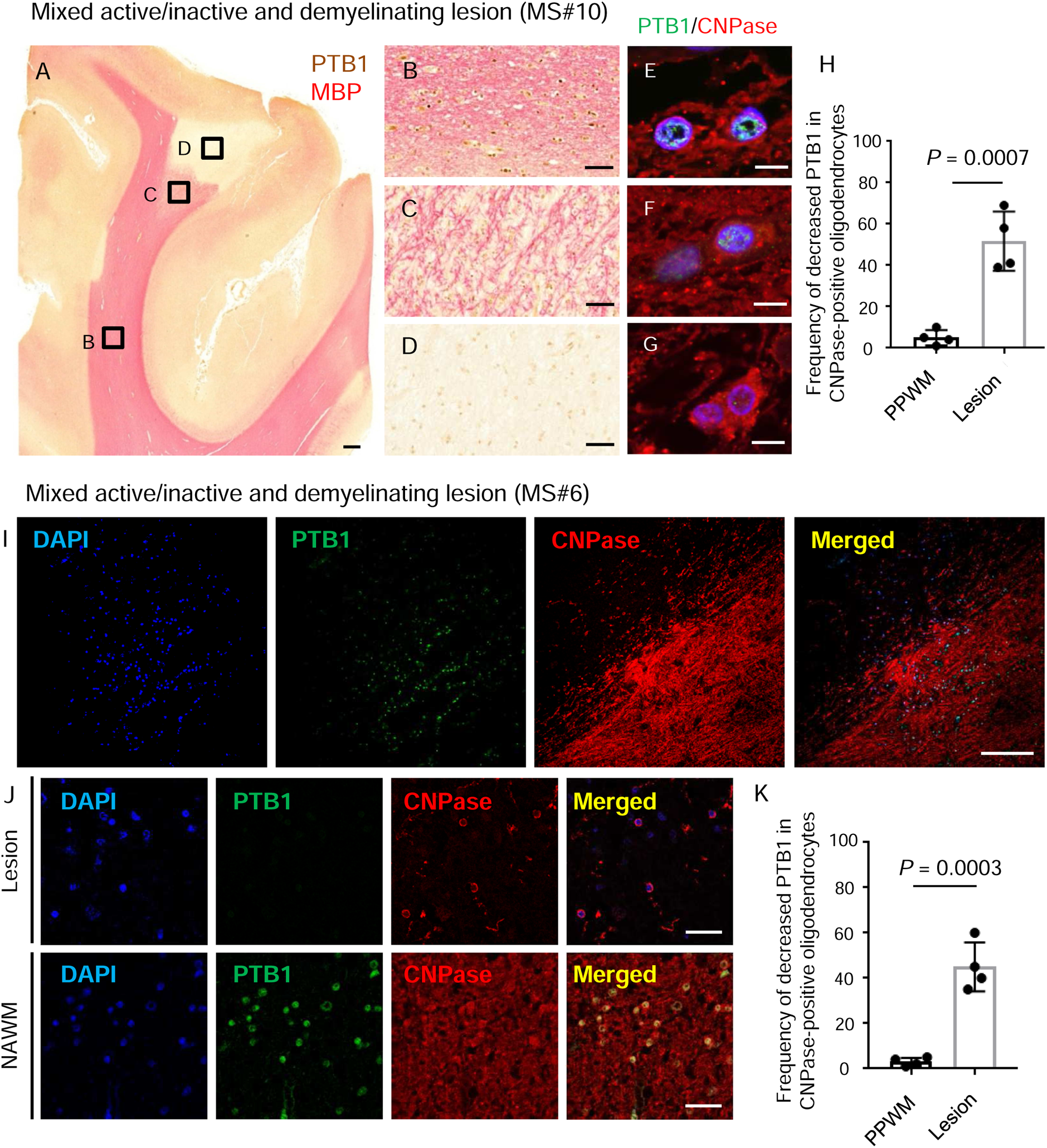
Reduced expression of PTB1 in oligodendrocytes of demyelinating lesions. (A-H) Mixed active/inactive and demyelinating subcortical white matter lesion from MS#10. (A-D) Double immunostaining for PTB1 (*brown*) and MBP (*pink*). (A) Macroscopic view of demyelinating subcortical lesion. Immunoreactivity for MBP is sharply demarcated in a subcortical area that has no detectable PTB1. (B) Immunoreactivity for PTB1 is present in the nucleus of glial cells, including oligodendrocytes, in NAWM. (C) Nuclear expression of PTB1 is diminished in an incompletely remyelinated area. (D) Expression of PTB1 is markedly decreased in glial cell nuclei in the demyelinated lesion. (E-G) Double immunofluorescence for PTB1 and CNPase. While immunoreactivity for PTB1 is preserved in the nuclei of CNPase-positive oligodendrocytes in NAWM (E), nuclear PTB1 is decreased in the remyelinated area (F) and markedly depleted in oligodendrocytes in the lesion center (G). (H) Frequency of decreased PTB1 expression in CNPase-positive oligodendrocytes. Decreased PTB1 in oligodendrocytes is significantly frequent in demyelinating lesions than PPWM. (I-K) Mixed active/inactive and demyelinating white matter lesion from MS#6 (I) Double immunofluorescence shows that nuclear expression of PTB1 is markedly diminished in the demyelinating lesion. (J) Higher magnification of PTB1 expression in the demyelinating lesion and normal-appearing white matter (NAWM). Although PTB1 is localized in nuclei of CNPase-positive oligodendrocytes of NAWM, PTB1 is decreased in CNPase-positive oligodendrocytes of the demyelinating lesion. (K) Decreased PTB1 expression in oligodendrocytes in active-inactive demyelinating lesions is significantly more frequent than in PPWM. Scale bars: 1 mm (A), 100 µm (I), 50 µm (B-D), 20 µm (J) and 10 µm (E-G).

### Alteration of RBPs in cortical plaques

Leukocortical mixed active/inactive demyelinating lesions from patients MS#2, 8, 10-12 had mild to moderate diminution of PTB2 expression in neurons within the demyelinated area compared to neurons in adjacent normal-appearing gray matter (NAGM) (MS#2 - Fig. 4A-H, MS#10 – Fig. 4I-L) (Table e-1). Although the expression of PTB2 was decreased in the nucleus in these cells, there was no evidence of cytoplasmic mislocalization or aggregate formation of PTB2 (Fig. 4E-G). The decrease in PTB2 expression in cortical neurons in leukocortical plaques in the case of MS#2 and #10 was statistically significant compared to cortical neurons in the PPGM (P < 0.0001) (Fig. 4H, L). In contrast to these findings, the expression of TDP-43 and FUS in neurons in leukocortical plaques, and of TDP-43, FUS and PTB2 in neurons of intracortical and subpial plaques had a normal expression in the nuclei and did not differ from that in NAGM (Table e-1).

**Figure 4.**
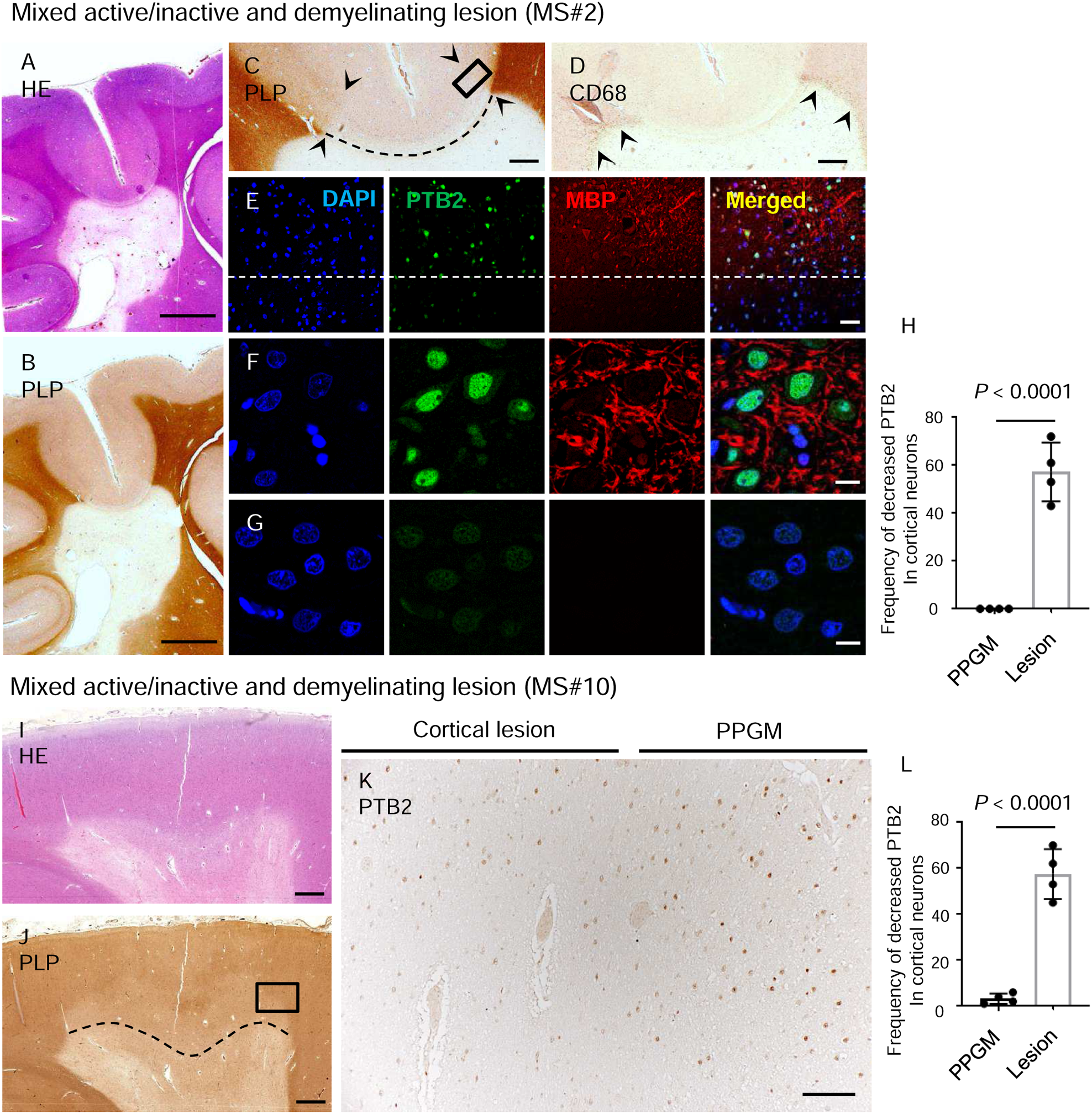
Reduced expression of PTB2 in neurons of cortical demyelinating lesions. (A, B) Serial sections of a mixed active/inactive demyelinating lesion. (A) HE stain shows hypocellularity, and (B) immunostaining for PLP shows sharply demarcated periventricular demyelination including (C) cortical demyelination (*arrowheads*), which is above the dotted line. (D) CD68-positive macrophages are restricted to the periphery of the lesion (*arrowheads*). (E-G) Higher magnification of double immunofluorescence of *square* area (at edge of cortical demyelination) in Panel C. (E) PTB2 has normal expression in the nuclei of neurons in the non-affected area of the gray matter (above the dashed line) (F). In contrast, the expression of PTB2 is markedly decreased in neurons in the demyelinated area of the gray matter (below the dashed line) (G). (H) Frequency of decreased PTB2 expression in cortical neurons. Decreased PTB2 in cortical neurons is significantly more frequent in cortical demyelinating lesions than PPGM. (I-L) Leukocortical mixed active/inactive demyelinating lesion MS#10. (I) HE stain shows subcortical white matter lesion. (J) The cortex above the dashed line and within the lesion is demyelinated. (K) A higher magnification of the region within the *rectangle* in panel J includes the boundary of cortical demyelination. PTB2 expression is diminished in the demyelinated region of the cortex, while PTB2 is preserved in the cortical neurons in same layer of PPGM. (L) Frequency of decreased PTB2 expression in cortical neurons. Decreased PTB2 in cortical neurons is significantly more frequent in cortical demyelinating lesions than PPGM. Scale bars: 5 mm (A, B), 1 mm (C, D, I, J), 100 µm (K), 50 µm (E) and 10 µm (F, G).

### TDP-43 proteinopathy in *in vitro* cultured primary human oligodendrocytes

We cultured primary human oligodendrocytes in a low glucose medium in order to model metabolic stress conditions thought to occur in MS lesions ^12, 13^. As shown in Fig. 5A, there is only a low level of cell death under N1 conditions at day 2 in culture. Levels modestly increase under LG conditions, as previously described ^12, 13^. By day 6, however, there was a significant increase in cell death under LG conditions. The percent of O4 cells showing predominantly nuclear TDP-43 staining was significantly reduced in the LG-treated cultures compared to the N1 counterparts (mean 53% for LG vs 82% for N1; P=0.015, Fig. 5B and illustrated in Fig. 5C). Nuclear depletion was observed in virtually all PI+ cells, as shown with the PI+ cells in Fig. 5C. Of note, the percent of PI-cells with nuclear depletion was greater in LG cultures versus N1 cultures (mean 56% for LG vs 86% for N1; P = 0.014, n=3, data not shown).

**Figure 5.**
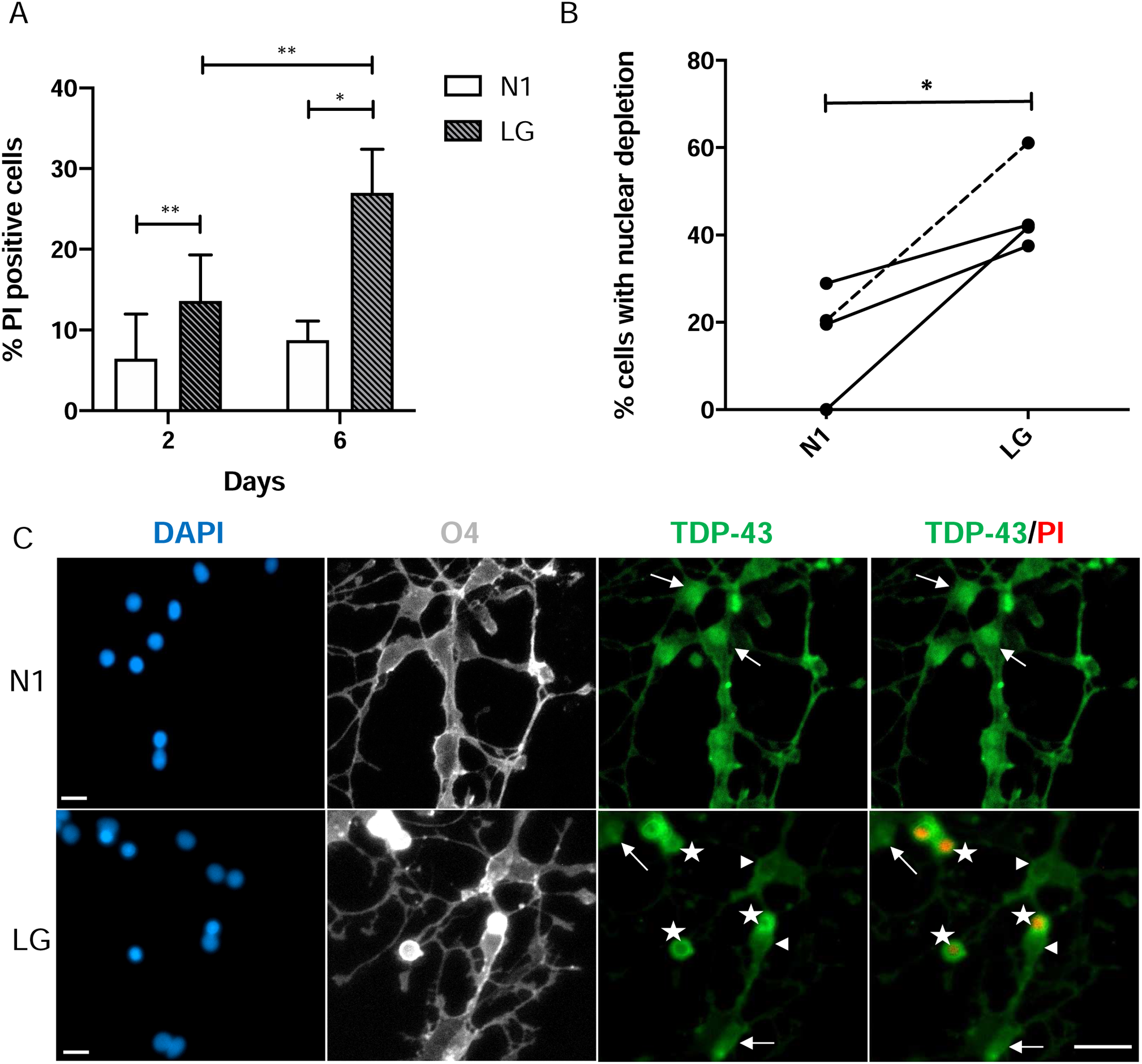
Nuclear depletion of TDP-43 in human primary OLs cultured in vitro under LG conditions. Data were derived from 3 adult cases and 1 pediatric case with no history of MS. (A) Cell death of human OLs was assessed using propidium iodide (PI) staining at 2 and 6 days under optimal (N1) and metabolic stress (LG- low glucose) culture conditions. (B) Depletion of nuclear expression of TDP-43 in cutures of oligodendrocytes after 2 days of treatment with LG compared to N1 condition. Solid lines (adult cases) and dashed line (pediatric case) connect cultures of oligodendrocytes obtained from the same biopsy tissue. (C) Representative images showing immunostained oligodendrocytes 2 days under N1 and LG conditions. From left side: DAPI (blue), O4 (gray), TDP-43 (green), and TDP-43 merged with PI (green and red respectively). Arrows show cells with nuclear expression of TDP-43; arrow heads show TDP-43 nuclear-depleted PI- cells; stars show TDP-43 nuclear-depleted PI+ cells. Scale bar = 10 µm. *P < 0.05, **P < 0.01.

## Discussion

Abnormalities of expression and localization of RBPs have been described in a number of diseases, including ALS, Huntington’s disease, and viral infections ^2^. In the present study, we focused on TDP-43, FUS, and PTB because these RBPs have an important impact on RNA biology and also because their mislocalization is thought to influence the pathogenesis of ALS and TMEV infections.

TDP-43 is a ubiquitously expressed RBP that predominantly resides in the nucleus, but shuttles across the nuclear membrane in association with mRNAs ^15^. A hallmark of almost all cases of ALS is disruption of nucleocytoplasmic trafficking with resultant nuclear depletion, cytoplasmic mislocalization, aggregation, cleavage, and phosphorylation of TDP-43 in neural cells ^16-18^. The decreased expression and mislocalization or TDP-43 are thought to cause abnormalities of splicing and RNA metabolism as well as add to nucleocytoplasmic transport disruption, thereby contributing to ALS pathogenesis ^19-22^. It is likely that cytoplasmic mislocalization of other RBPs in addition to TDP-43 adds to the cellular dysfunction in ALS ^23^.

In the present study, we demonstrate a number of abnormalities in expression and localization of RBPs in MS lesions and in *in vitro* cultured oligodendrocytes. We found that TDP-43 was mislocalized in oligodendrocytes in demyelinated lesions in MS, as was the case in TMEV infections. Of note, TDP-43 is known to bind to 100s of mRNAs, including mRNAs encoding PLP, myelin basic protein, myelin oligodendrocyte glycoprotein, and myelin-associated glycoprotein, and to play a key role in RNA metabolism and splicing ^24^. Importantly and relevant to our findings is a recent report that an experimental decrease in expression of TDP-43 in mature oligodendrocytes in mice leads to demyelination and RIPK1-mediated necroptosis of oligodendrocytes ^6^; of note, necroptosis has been reported to occur in MS as well as experimental models of MS ^25^. A conclusion of that investigation was that the TDP-43 knockdown led to a reduction in myelin gene expression, and that TDP-43 is indispensable for oligodendrocyte survival and demyelination ^6^. These findings suggest that nuclear depletion and mislocalization of TDP-43 in MS lesions would similarly lead to demyelination and, in some cases, death of oligodendrocytes.

We found a decrease in PTB1 in oligodendrocytes in mixed active/inactive demyelinating lesions, as well as a decrease in PTB2 in neurons in cortical plaques. PTB1 and PTB2 are paralogous RBPs that are encoded by related genes ^7^. PTB1 is not expressed in mature neurons and muscle, while PTB2 is expressed in these cells and others. These RBPs function in regulating alternative splicing, and also play a role in translation, mRNA stability, and polyadenylation. The control of splicing is especially important in the CNS because of the myriad of mRNA isoforms that have key roles in development and function. Splicing in oligodendrocytes and neurons in MS demyelinated regions is likely impacted by the nuclear depletion and cytoplasmic mislocalization of PTB. Importantly, PTB is involved in the differentiation of neural precursor cells ^7, 8^. In this way, PTB2 nuclear depletion and cytoplasmic mislocalization in neurons of cortical plaques may contribute to neurodegeneration and the cognitive decline associated with MS.

In summary, we found that there is disruption of TDP-43 and PTB expression and localization that varies in different neural cell types in MS plaques. This variation may have resulted from differences in the protein composition of the nuclear pore complex in different cell types ^26^. Furthermore, different subtypes of MS lesions may manifest continuing changes of RBP abnormalities over time because of the dynamic nature of demyelinating lesions and the varying inflammatory milieu. It may be that this changeable and very dynamic nature of MS lesions is the reason that active plaques from MS#14 had a normal localization of TDP-43. Also of importance is the fact that MS is a heterogeneous disease – and therefore it is not surprising that forms of MS that are different from classical and typical cases may not share the same RBP abnormalities seen in more prototypic cases of MS. Perhaps this was the reason that an active plaque from a biopsy from a tumefactive lesion of MS#4 (in a patient who had only one additional clinical problem over decades of observation) had a normal localization of TDP-43 (Table e-1).

The nucleocytoplasmic transport abnormalities in MS that we identified may have resulted from a number of possible causes. Probably most relevant are reports that inflammation can lead to mislocalization of proteins in neural cells. Correia et al. ^27^ found that mislocalization of TDP-43 occurred in: cultured microglia and astrocytes following exposure to lipopolysaccharide (LPS); motor neuron-like NSC-34 cells after treatment with tumor necrosis factor-alpha (TNFa); and motor neurons of mutant TDP-43 transgenic mice following LPS intraperitoneal injections. Kim et al. ^28^ reported that treating neuronal cultures with glutamate and TNFa led to mislocalization of HDAC1 with resultant axonal damage. The latter investigators also detected abnormal cytoplasmic localization of HDAC1 in damaged axons in MS patients and in mice with cuprizone-induced demyelination. Salapa et al. ^29^ found that interferon-γ led to cytoplasmic mislocalization of heterogeneous ribonuclear protein (hnRNP) A1, an RBP. These investigators also reported that neurons in a region of an MS brain (in which no pathology was described) had nuclear depletion and cytoplasmic mislocalization of hnRNP A1, which was aggregated in stress granules.

Our in vitro culture conditions were selected to model metabolic stress conditions that are thought to occur in MS lesions ^12, 13^. Antel and colleagues previously showed that these conditions were associated with an initial withdrawal of cell processes, modelling the dying-back of oligodendrocyte processes observed in MS lesions (as well as cuprizone-induced demyelination ^30^ and TMEV-induced demyelination ^31^. These changes were reversible if culture conditions were restored within a subsequent 48 hours. If continued past this time, however, significant cell death occurs by 6 days, as shown in Fig. 5A, with activation of an autophagy response. In the current study, we found TDP-43 nuclear depletion was increased under LG conditions after 2 days in culture, a time when cell death levels as detected by PI staining were low. Importantly, we specifically observed nuclear depletion in cells that were still PI negative, in addition to PI+ cells that also showed nuclear depletion of TDP-43. These results are consistent with in situ data showing nuclear depletion of TDP-43 in oligodendrocytes with intact oligodendrocyte cell bodies. Of note, TNFa also leads to dying back of cultured oligodendrocytes, although in this case it was observed in newborn rat-derived oligodendrocytes ^12^. The combined in vitro and in situ results suggest that the TDP-43 nuclear depletion reflects a cellular stress response that could be mediated both by the metabolic conditions and inflammatory mediators of MS lesions. Of note, no difference in TDP-43 transcripts was found in a microarray data set derived from oligodendrocytes under N1 vs LG conditions for 2 days ^12^, suggesting that any change in TDP-43 protein levels is a result of translational regulation, presumably from stress, such as from low glucose ^32^ or inflammatory factors, triggering the integrated stress response. Activation of the integrated stress response has been previously implicated in the pathogenesis of MS ^33^.

Our results suggest that correcting the expression and localization of RBPs in MS may ameliorate disease. In addition, this direction may lead to normal localization of key transcription factors and proteins that are required for efficient myelination and remyelination in oligodendrocytes and oligodendrocyte precursor cells ^34-37^. Importantly, nuclear export inhibitors have been found to attenuate myelin oligodendrocyte glycoprotein-induced experimental autoimmune encephalomyelitis (as well as kainic acid-induced axonal damage) by limiting areas of myelin damage, preserving myelinated and unmyelinated axon integrity, and decreasing inflammation ^38^. Nuclear export inhibitors have also been found to attenuate disease and to be neuroprotective in experimental models of ALS ^39^, including a mutant TDP-43 mouse model ^19^, and Huntington’s disease ^40^ (which, like ALS, has abnormalities of nucleocytoplasmic transport). Furthermore, nucleocytoplasmic transport is being targeted in patients with cancer in addition to neurological diseases – and a clinical trial with a nuclear export inhibitor is in progress in ALS.

Altered nucleocytoplasmic transport leading to abnormal expression and mislocalization of RBPs and other macromolecules may not only contribute to the demyelination and neurodegeneration in MS, but also underlie a number of other disease states, both infectious as well as non-infectious. The availability of drugs that target nucleocytoplasmic transport may provide new and novel treatment possibilities for these disorders.

## Acknowledgments

We thank the UCLA Human Brain and Spinal Fluid Resource Center and the Rocky Mountain MS Center Tissue Bank (supported in part by a grant from the National Multiple Sclerosis Society) for specimens from autopsied MS cases.

## Appendix

Authors

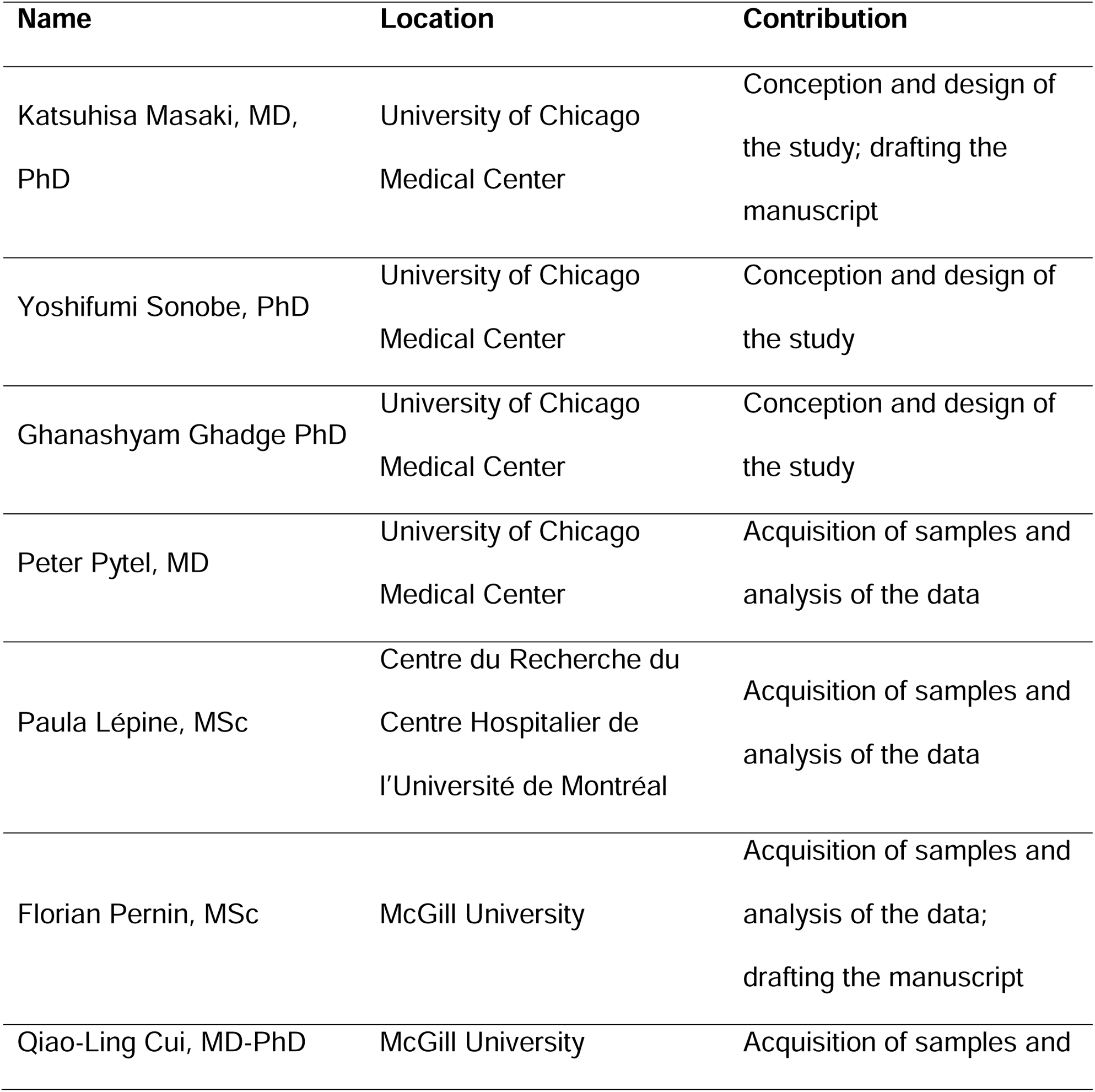
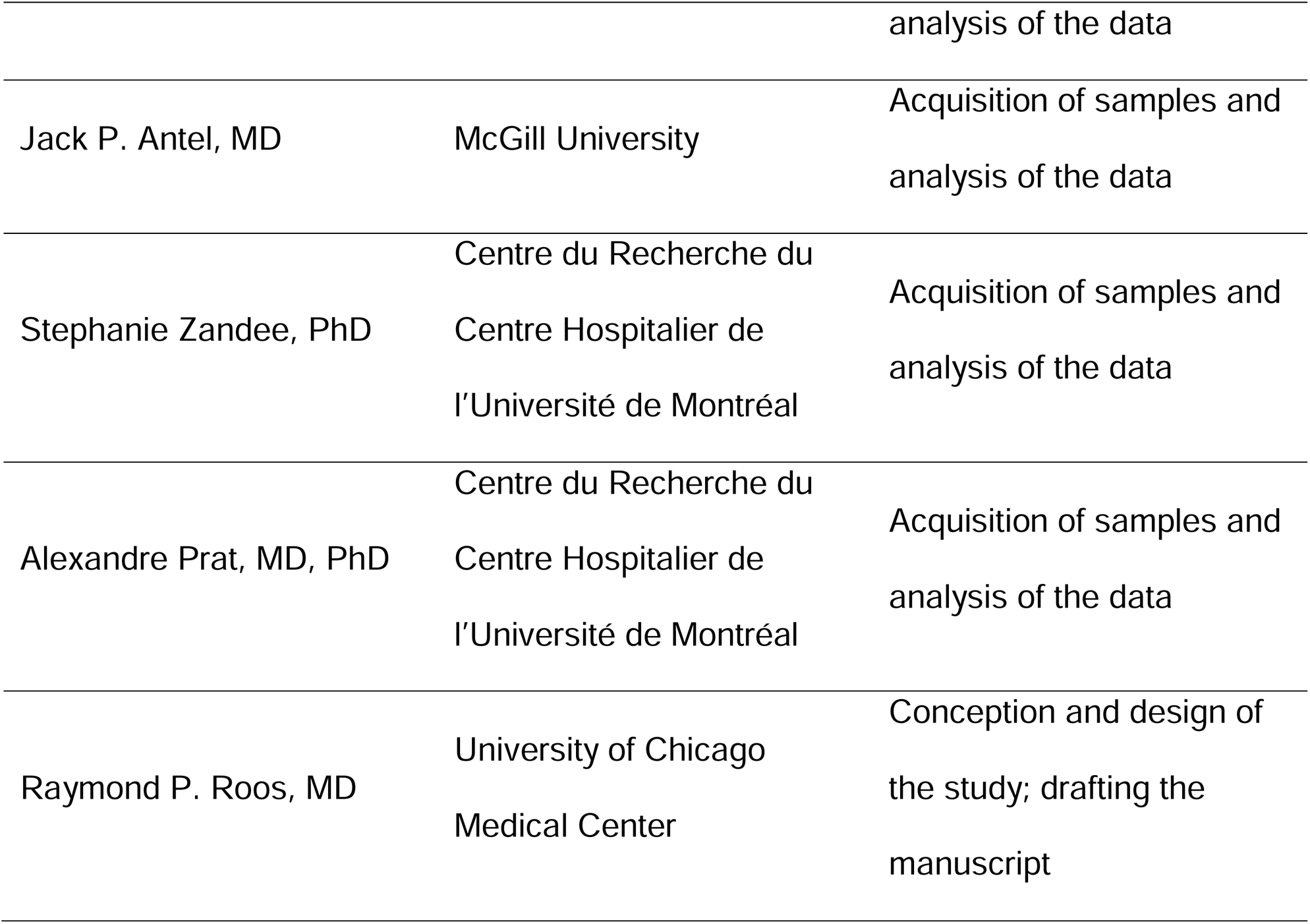

## Supplemental Data

**Figure e-1: TDP-43 mislocalization in oligodendrocytes of active demyelinating lesion of MS.**

(A) Double immunofluorescence for TDP-43 and CNPase in inactive demyelinating lesion from MS#6. The dotted line shows the boundary of demyelinating lesion and periplaque white matter (PPWM). Nuclear expression of TDP-43 is seen in the demyelinating lesion as well as PPWM. (B-D) Double immunofluorescence for TDP-43 and CNPase in active demyelinating lesion from MS#13. (B) Nuclear expression of TDP-43 is seen in PPWM, while nuclear staining of TDP-43 is markedly diminished in the demyelinating lesion. (C) Higher magnification of double immunofluorescence. Nuclear TDP-43 is depleted in CNPase-positive oligodendrocyte of the active demyelinating lesion, whereas TDP-43 is expressed in the nuclei of oligodendrocytes of PPWM. (D) Frequency of TDP-43 mislocalization in CNPase-positive oligodendrocytes. TDP-43 mislocalization in oligodendrocytes is significantly higher in demyelinating lesions than PPWM. Scale bars: 100 µm (A, B) and 10 µm (C).

**Table e-1.** MS cases.

| Patient # | Age | Sex | Clinical subtype* | White matter lesion**<br>(number examined) | Oligodendrocyte |  | Gray matter lesion†<br>(number examined) | Neuron<br>Decreased nuclear PTB2 |
| --- | --- | --- | --- | --- | --- | --- | --- | --- |
|  |  |  |  |  | TDP-43<br>Proteinopathy† | Decreased nuclear PTB1 |  |  |
| 1 | 38 | F | MS | I (1) | – | – | None | – |
| 2 | 66 | F | MS | A/I (4) | – | ++ | LC (1) | + |
| 3 | 53 | F | MS | A (1) | ++ | – | None | – |
|  |  |  |  | A/I (1) | – | – |  |  |
| 4<sup>s</sup> | 30 | F | T | A (1) | – | – | None | – |
| 5 | 56 | M | Prog | A/I (1) | – | – | None | – |
|  |  |  |  | I (3) | – | – |  |  |
| 6 | 63 | M | Prog | A/I (3) | – | ++ | None | – |
| 7 | 47 | F | Prog | I (3) | – | – | None | – |
| 8 | 38 | M | MS | A/I (6) | – | – | SP (1) | – |
|  |  |  |  |  |  |  | IC (2) | – |
|  |  |  |  |  |  |  | LC (2) | + |
| 9 | 42 | F | MS | A/I (2) | – | – | SP (4) | – |
|  |  |  |  |  |  |  | IC (2) | – |
| 10 | 30 | M | RRMS | A/I (6) | – | +++ | SP (2) | – |
|  |  |  |  | I (1) | – | – | LC (3) | ++ |
| 11 | 47 | M | Prog | A/I (2) | – | ++ | SP (3) | – |
|  |  |  |  |  |  |  | IC (1) | – |
|  |  |  |  |  |  |  | LC (1) | + |
| 12 | 42 | F | Prog | A/I (1) | – | ++ | SP (4) | – |
|  |  |  |  | I (1) | – | – | LC (1) | ++ |
| 13 | 60 | F | Prog | A (3) | ++ | + | None | – |
| 14 | 26 | M | RRMS | A (3) | – | – | None | – |
|  |  |  |  | I (1) | – | – |  |  |

\*A: active lesion, A/I: mixed active/inactive lesion, I: inactive lesion.
<sup>†</sup>TDP-43 mislocalization and/or decreased nuclear expression.
<sup>‡</sup>IC: intracortical lesion, LC: leukocortical lesion, SP: subpial lesion.
<sup>§</sup>MS#4: tissue was a biopsy from a tumefactive lesion.
<sup>\*\*</sup>MS: multiple sclerosis with unclear clinical classification of the case, Prog: Progressive multiple sclerosis, RRMS: relapse-remitting multiple sclerosis. T: tumefactive

**Table e-2.** Antibodies used for immunohistochemistry and immunofluorescence

| Antigen |  | Type | Dilution | Source |
| --- | --- | --- | --- | --- |
| RNA-binding protein | TDP-43 (N-terminal) | Rabbit | 1:1000 | Proteintech, Rosemont, IL, USA |
|  | TDP-43 (1D3, phospho-S409/410) | Rat | 1:100 | EMD Millipore, Burlington, MA, USA |
|  | PTB1 | Rabbit | 1:100 | Abcam, Cambridge, UK |
|  | PTB2 | Rabbit | 1:100 | Abcam |
|  | FUS (11570-1-AP) | Rabbit | 1:50 | Proteintech |
| Oligodendrocyte/myelin | PLP | Rabbit | 1:400 | Sigma Aldrich |
|  | MBP (F-6) | Mouse | 1:100 | Santa Cruz Biotechnology, Dallas, Tx, USA |
|  | CNPase | Mouse | 1:100 | Abcam |
| Macrophage | CD68 (KP1) | Mouse | 1:300 | Abcam |

Formatted: Font: (Default) +Headings (Arial)
Formatted Table
Formatted: Font: (Default) +Headings (Arial), German (Germany)

