## Supplement methods and figures for "RNA-binding protein altered expression and mislocalization in multiple sclerosis"

**Ethics Statement**

The study involved tissue from human subjects. The study was approved by The University of Chicago Institutional Review Board for Clinical Research. Informed written consent for an autopsy at the University of Chicago was obtained from an immediate member of the deceased’s family. The autopsies on patients from the Centre de Recherche du Centre Hospitalier de l’Université de Montréal had informed consent and was in accordance with institutional guidelines and approval by the local Centre Hospitalier de l’Université de Montréal ethics committee (HD04.046 and BH07.001). The use of tissue from the Montreal Neurological Institute, McGill University McGill University Health Center Research Ethics Board was approved by the McGill University Health Center Research Ethics Board.

**Human Samples**

Immunohistochemical studies were performed on autopsy brain specimens from 14 patients with diagnoses that included MS, ALS, and control (myasthenia gravis) on the basis of the clinical syndrome and neuropathological findings[^1^](https://mc.manuscriptcentral.com/actn?DOWNLOAD=TRUE&PARAMS=xik_2r3jcHwbhQWagRtgtKqivxVQVMKoatmwQgtVx6hknhfz2GUjUMCcUpZFiVUebU3do6M1HkvhghMXqgW8BeJLJNLRYts5wRzZ25QWCXmwVAL2ov4kaEEKzpEuxtGgdYbsmmUZbQs5gfo96dwUfXVk6VLVS3YaE2h57JoBcgEaAJsNhzEZsV5kzuq1NtmcmBWcPL7pLQ6Gpqk5qx9Lyy6TEJCnFT8Ck6oDVFKuNLeF5nc2x56Tp#_ENREF_1)^,^ [^9^](https://mc.manuscriptcentral.com/actn?DOWNLOAD=TRUE&PARAMS=xik_2r3jcHwbhQWagRtgtKqivxVQVMKoatmwQgtVx6hknhfz2GUjUMCcUpZFiVUebU3do6M1HkvhghMXqgW8BeJLJNLRYts5wRzZ25QWCXmwVAL2ov4kaEEKzpEuxtGgdYbsmmUZbQs5gfo96dwUfXVk6VLVS3YaE2h57JoBcgEaAJsNhzEZsV5kzuq1NtmcmBWcPL7pLQ6Gpqk5qx9Lyy6TEJCnFT8Ck6oDVFKuNLeF5nc2x56Tp#_ENREF_9) (Table S1). Four cases (MS #1-4) of MS, three cases of ALS, two cases of cerebral infarction, and one case of myasthenia gravis were obtained from the Department of Pathology, University of Chicago. Additional cases of MS were provided by the Human Brain and Spinal Fluid Resource Center (Los Angeles, CA) (MS #5-7), Rocky Mountain MS Center Tissue Bank (Westminster, CO) (MS #8-10), and the Neuroimmunology Research Laboratory, Department of Neurosciences, Faculty of Medicine, Centre de Recherche du Centre Hospitalier de l’Université de Montréal (MS #11-14). The latter autopsy samples were preserved and lesions classified using Luxol Fast Blue/hematoxylin and eosin staining, as previously published [^10^](https://mc.manuscriptcentral.com/actn?DOWNLOAD=TRUE&PARAMS=xik_2r3jcHwbhQWagRtgtKqivxVQVMKoatmwQgtVx6hknhfz2GUjUMCcUpZFiVUebU3do6M1HkvhghMXqgW8BeJLJNLRYts5wRzZ25QWCXmwVAL2ov4kaEEKzpEuxtGgdYbsmmUZbQs5gfo96dwUfXVk6VLVS3YaE2h57JoBcgEaAJsNhzEZsV5kzuq1NtmcmBWcPL7pLQ6Gpqk5qx9Lyy6TEJCnFT8Ck6oDVFKuNLeF5nc2x56Tp#_ENREF_10)

**Tissue preparation and immunohistochemistry/immunofluorescence**

Brain tissue samples were fixed in 10% buffered formalin and processed into paraffin sections (5-µm thick). The sections were routinely subjected to hematoxylin and eosin (HE) or processed for immunohistochemistry or immunofluorescence. For the latter processing, sections were deparaffinized in xylene and rehydrated through an ethanol gradient. Deparaffinized sections were then incubated with 0.3% hydrogen peroxide in absolute methanol for 30 min at room temperature to inhibit endogenous peroxidase, and then overlaid overnight at 4°C with primary antibodies. The primary antibodies used for immunohistochemistry and immunofluorescence are listed in Table S2. The following markers were used for specific neural cells: astrocytes - glial fibrillary acidic protein (GFAP); myelin/oligodendrocytes - myelin basic protein (MBP), proteolipid protein (PLP), and 2',3'-Cyclic-nucleotide 3'-phosphodiesterase (CNPase); macrophages - CD68.

For immunohistochemical studies, sections were rinsed after the primary antibody overlay and then labeled using an enhanced indirect immunoperoxidase method, with the reaction product developed using a solution of 3, 3’-diaminobenzidine, and counterstained with hematoxylin. Double immunostaining was carried out with two enzyme systems, peroxidase and alkaline phosphatase, followed by staining with Vector Red (Vector Laboratories).

For immunofluorescence studies, sections were rinsed after the primary antibody overlay and incubated with either Alexa 488-conjugated goat anti-rabbit IgG or Alexa 594-conjugated goat anti-mouse IgG (Invitrogen), and counterstained with DAPI. Images were captured using a confocal laser microscope system (Leica TCS SP5, Leica Microsystems, Wetzlar, Germany). A sequential multiple fluorescence scanning mode was used to avoid nonspecific overlap of signals. A 0.1% Sudan Black B solution was used to reduce lipofuscin autofluorescence.

**Supporting information**

**Figure S1: TDP-43 expression in inactive demyelinating lesion of MS.**

Double immunofluorescence for TDP-43 and CNPase in inactive demyelinating lesion from MS#6. The dotted line shows the boundary of demyelinating lesion and periplaque white matter (PPWM). Nuclear expression of TDP-43 is seen in the demyelinating lesion as well as PPWM. Scale bar: 100 µm.

**Figure S2: TDP-43 mislocalization in oligodendrocytes of active demyelinating lesion of MS.**

(A-C) Double immunofluorescence for TDP-43 and CNPase in active demyelinating lesion from MS#13. (A) Nuclear expression of TDP-43 is seen in PPWM, while nuclear staining of TDP-43 is markedly diminished in the demyelinating lesion. (B) Higher magnification of double immunofluorescence. Nuclear TDP-43 is depleted in CNPase-positive oligodendrocyte of the active demyelinating lesion, whereas TDP-43 is expressed in the nuclei of oligodendrocytes of PPWM. (C) Frequency of TDP-43 mislocalization in CNPase-positive oligodendrocytes. TDP-43 mislocalization in oligodendrocytes is significantly higher in demyelinating lesions than PPWM. Scale bars: 100 µm (A) and 10 µm (B).

Figure. S1

Mixed active/inactive and demyelinating lesion (MS#6)

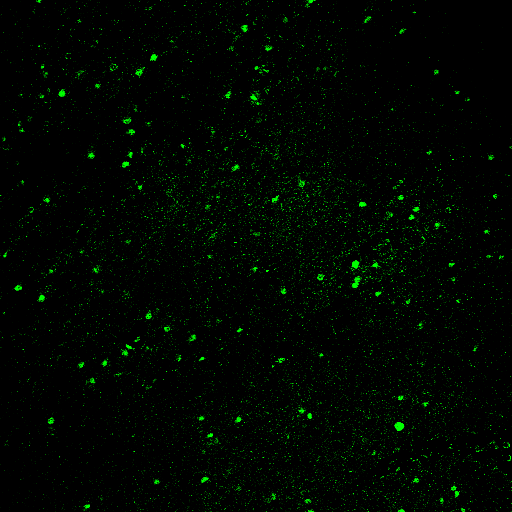

**TDP-43**

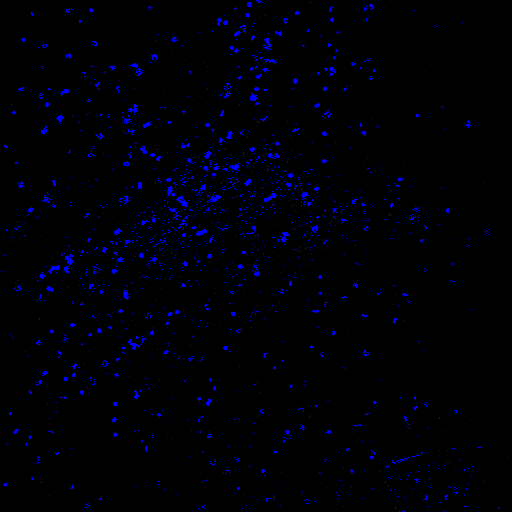

**DAPI**

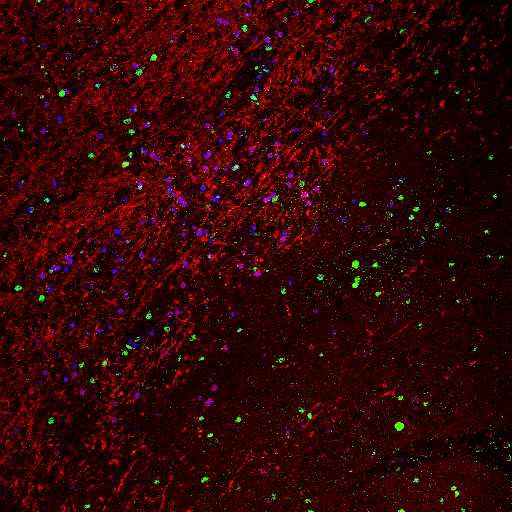

**Merged**

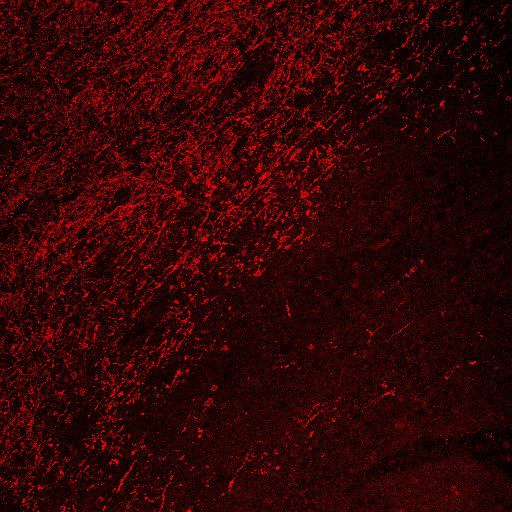

**CNPase**

**Inactive lesion**

Figure. S2

Active and demyelinating lesion (MS#13)

PPWM	Lesion

A

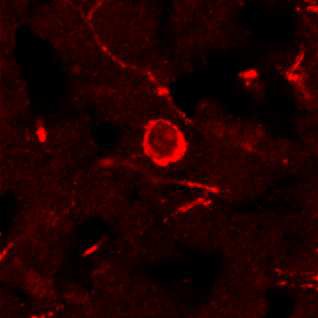

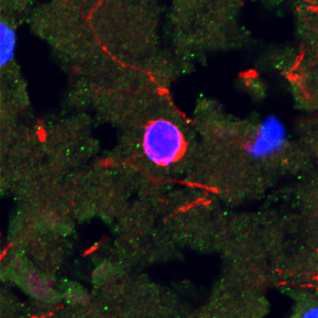

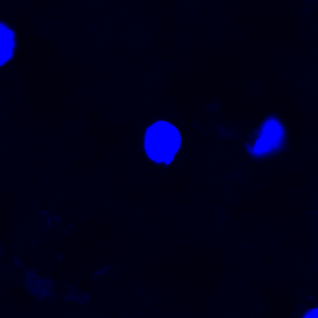

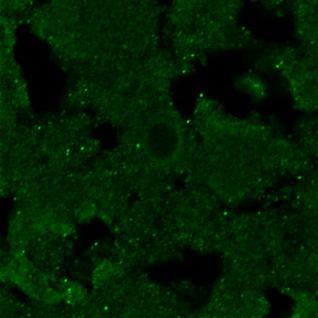

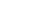

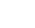

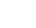

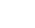

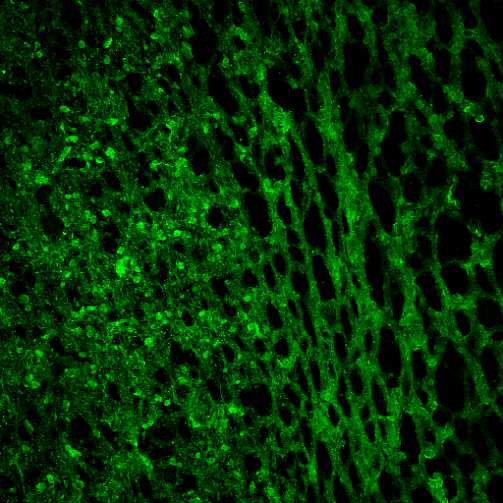

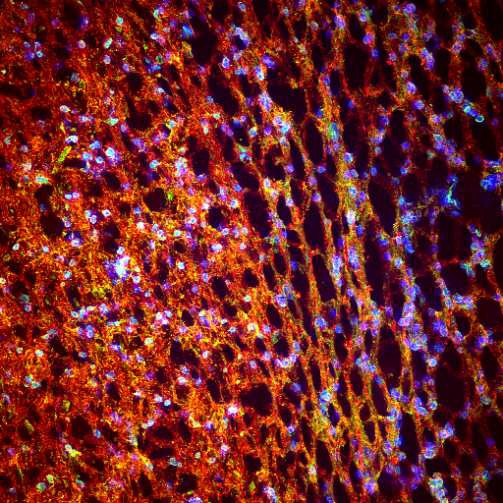

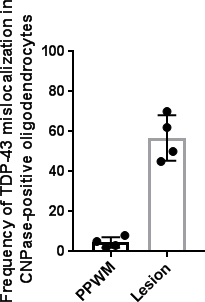

**TDP-43**

C

**Merged**

100

*P* = 0.0001

80

60

40

20

0

**DAPI**

**TDP-43**

**CNPase**

**Merged**

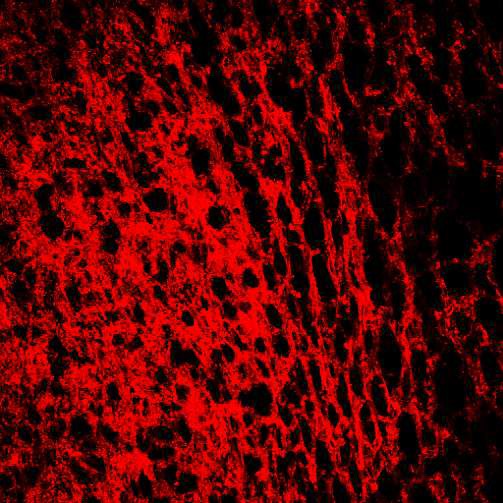

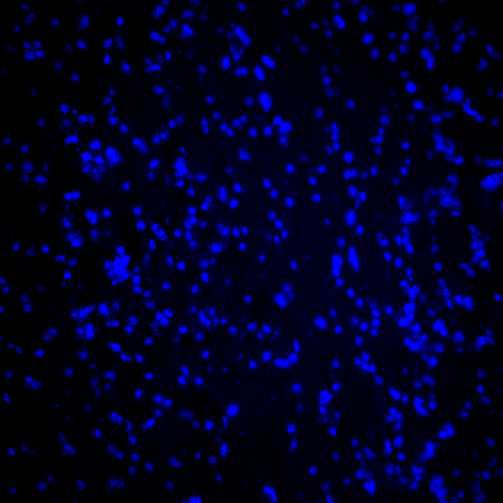

**DAPI**

**CNPase**

B

Active lesion

Frequency of TDP-43 mislocalization in CNPase-positive oligodendrocytes

PPWM

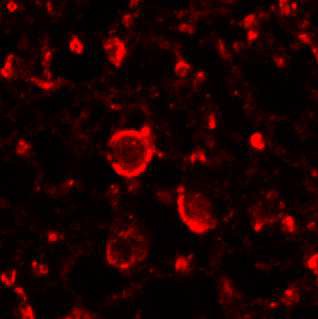

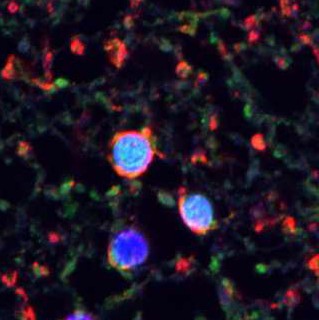

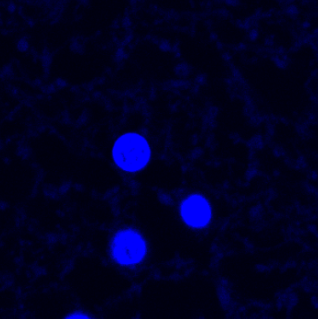

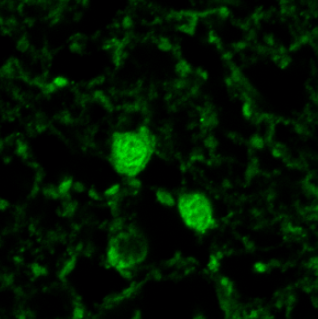

**DAPI**

**TDP-43**

**CNPase**

**Merged**
